# L1CAM is not Associated with Extracellular Vesicles in Human Cerebrospinal Fluid or Plasma

**DOI:** 10.1101/2020.08.12.247833

**Authors:** Maia Norman, Dmitry Ter-Ovanesyan, Wendy Trieu, Roey Lazarovits, Emma J.K. Kowal, Ju Hyun Lee, Alice S. Chen-Plotkin, Aviv Regev, George M. Church, David R. Walt

## Abstract

Neuron-derived extracellular vesicles (NDEVs) present a tremendous opportunity to learn about the biochemistry of brain cells in living patients. L1CAM is a transmembrane protein expressed in neurons that is presumed to be found on NDEVs in human biofluids. Previous studies have used L1CAM immuno-isolation from human plasma to isolate NDEVs for neurodegenerative disease diagnostics. We developed a panel of ultrasensitive Single Molecule Array (Simoa) assays for known EV markers, as well as L1CAM, and applied it to study EVs in human plasma and cerebrospinal fluid (CSF). We fractionated plasma and CSF by size exclusion chromatography (SEC) and density gradient centrifugation (DGC) to separate EVs from free proteins. We observed that L1CAM did not elute in the EV fractions, but rather eluted in the free protein fractions. We found that L1CAM is present as a free protein in human plasma and CSF, possibly due to proteolytic cleavage and/or alternative splicing. We further demonstrate that the isoforms found in CSF and plasma are different. These data collectively establish that L1CAM in plasma is not EV associated and should therefore not be used for NDEV isolation. Importantly, the framework and tools described herein will allow for evaluation of other potential candidate markers for isolation of NDEVs.

## Introduction

Developing therapeutics for neurological and psychiatric conditions has lagged far behind other medical disciplines partly due to the inability to perform brain biopsies on living individuals. Our current understanding of brain diseases relies mainly upon postmortem tissue analysis after neurodegeneration and cell death have already occurred. Typically, this postmortem analysis is coupled with data gathered from GWAS studies, induced pluripotent stem cell (iPSC) disease models and animal models. Although these techniques have yielded many insights, fundamental questions about the underlying biochemical processes of neurological and psychiatric disease still remain. Having access to the proteomic and transcriptomic profiles of neurons and other brain cells in living patients would be a great asset to our understanding of neuroscience. Furthermore, the ability to measure biochemical changes in response to medication would be a novel and useful tool for drug development.

One potential approach to learning about the living brain would be to analyze extracellular vesicles (EVs). EVs, which are secreted from many cell types and found in all biofluids, contain proteins and RNAs from their cell of origin [1]. There has been a great deal of excitement about capturing EVs secreted from neurons and characterizing their contents as a window into neurological processes. Over the past several years, a large number of studies have reported the immuno-capture of putatively neuron-derived EVs (NDEVs) and subsequent measurement of proteins and RNA implicated in neurodegenerative and psychiatric diseases. Almost all of these studies [2-46] used as a handle for EV capture the transmembrane protein L1CAM, a cell adhesion molecule implicated in neural development [47]. The choice of L1CAM relied on its known expression in neurons, presence in plasma, and the availability of high-affinity monoclonal commercial antibodies.

However, there are reasons that L1CAM may not be a good marker for NDEV immunocapture. First, L1CAM is expressed widely outside of the brain, including on non-neuronal cell types [48]. Furthermore, L1CAM has been shown to exist in soluble as well as transmembrane forms: an alternative splicing isoform which excludes the exon encoding the transmembrane domain has been observed [48] as well as various cleavage events by extracellular proteases that separate the extracellular domain (90% of the protein) from the membrane anchor [49, 50]. It is therefore not safe to assume that L1CAM present in extracellular fluids is necessarily vesicle-associated, but this must be directly validated.

Currently, publications that report measuring proteins inside L1CAM EVs use one of two antibodies to the L1CAM ectodomain (clones UJ127 or 5G3) [2-46]. However, if L1CAM is cleaved or alternatively spliced in the CSF and plasma, then it is possible that free soluble L1CAM is being captured, rather than EVs. Here we develop both a framework and the necessary tools for evaluating whether a given protein is a good candidate for neuron-specific EV pulldown. Using Simoa technology we analyze Density Gradient Centrifugation (DGC) and Size Exclusion Chromatography (SEC) fractions from CSF and plasma to demonstrate that L1CAM does not elute in the fractions that contain EVs; instead it elutes in the fractions where we observe free proteins. Furthermore, utilizing western blotting, mass spectrometry and RNA-Seq analysis, we demonstrate that CSF and plasma contain different isoforms of L1CAM. These data cast substantial doubt on the feasibility of using L1CAM to capture neuron-derived EVs.

## Results

In light of research demonstrating that both cleaved and alternatively spliced forms of L1CAM exist, we first sought to evaluate whether L1CAM in plasma and CSF is EV associated, or if it is a free protein. To do so, we fractionated biofluids using techniques for separating EVs from free proteins, including both DGC and SEC. We hypothesized that if L1CAM were predominantly associated with EVs, it would elute along with well-documented EV markers such as the tetraspanins CD9, CD63 and CD81 [1]. Alternatively, if L1CAM was predominantly a free protein, it would elute along with abundant free proteins such as albumin (schematically represented in Figure 1a). We first demonstrated that when purified L1CAM-expressing EVs from cell culture are put through SEC, they elute in earlier fractions but when recombinant L1CAM protein is put through SEC, it elutes in later fractions (Figure 1b). Although there is some overlap at the tails, these distributions are clearly distinct, and thus we refer to these samples as molecular standards for which fractions constitute vesicle-associated vs. free protein throughout the rest of this study.

**Figure 1:**
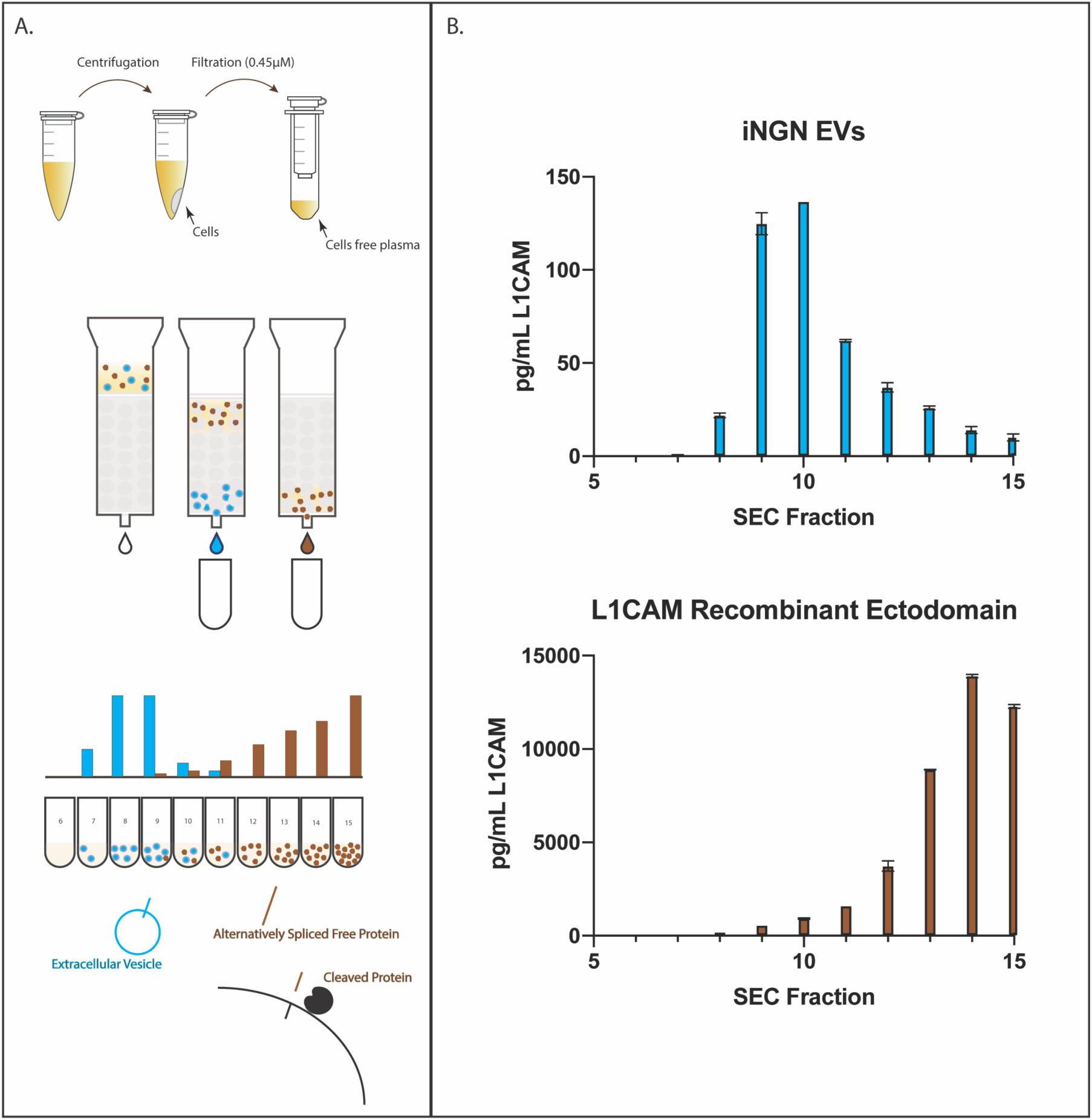
Method for evaluating whether L1CAM is EV associated or free in biofluids. A) Schematic representation of our method for evaluating whether L1CAM is EV associated using Size Exclusion Chromatography B) Size Exclusion Chromatography of cell culture EVs expressing L1CAM (top) and a free recombinant L1CAM protein (bottom)

Next, we sought to replicate this protocol with human biofluids, such as CSF and plasma. One technical challenge of working with human biofluids is that EVs are present in extremely low concentrations in these biofluids after fractionation. To ensure that we could quantify these low-abundance, putatively neuronally derived EVs, we developed and optimized high sensitivity assays for EV markers (CD9, CD63 and CD81) as well as for L1CAM and albumin (Supplementary Figure 1-2 & Supplementary Table 1). We utilized Simoa technology developed in our laboratory [51] to measure these low abundance targets in fractionated CSF and plasma.

### Evaluation of plasma and CSF by DGC as well as SEC

To evaluate whether L1CAM elutes in fractions with EVs or with free protein, we utilized two different techniques that separate EVs by size and density, respectively. The benefit of comparing the results of both techniques is that while DGC produces very pure EVs, it has relatively low yield; on the other hand, SEC can give high yields of EVs but is more likely to contain contaminating proteins [52]. The use of these two complementary methods allows for more accurate determination of whether L1CAM elutes as a free protein or is EV-associated.

When CSF and plasma were fractionated by SEC, the tetraspanins were detected primarily in fractions 7-9, while L1CAM was detected with a similar distribution to albumin in fractions 11-14 (Figure 2). When CSF and plasma were fractionated by DGC, we observed the same pattern, with the tetraspanin signal eluting in different fractions than the L1CAM and albumin signals: in CSF, the tetraspanins eluted in fractions 5-10, while albumin and L1CAM eluted in Fractions 1-5 (Figure 3a). In plasma, the tetraspanins eluted in fractions 7-11 while L1CAM, like albumin, eluted in fractions 1-6 (Figure 3b). Thus, with either of these two fractionation methods, L1CAM in human biofluids behaves as a free protein, not an EV associated protein.

**Figure 2:**
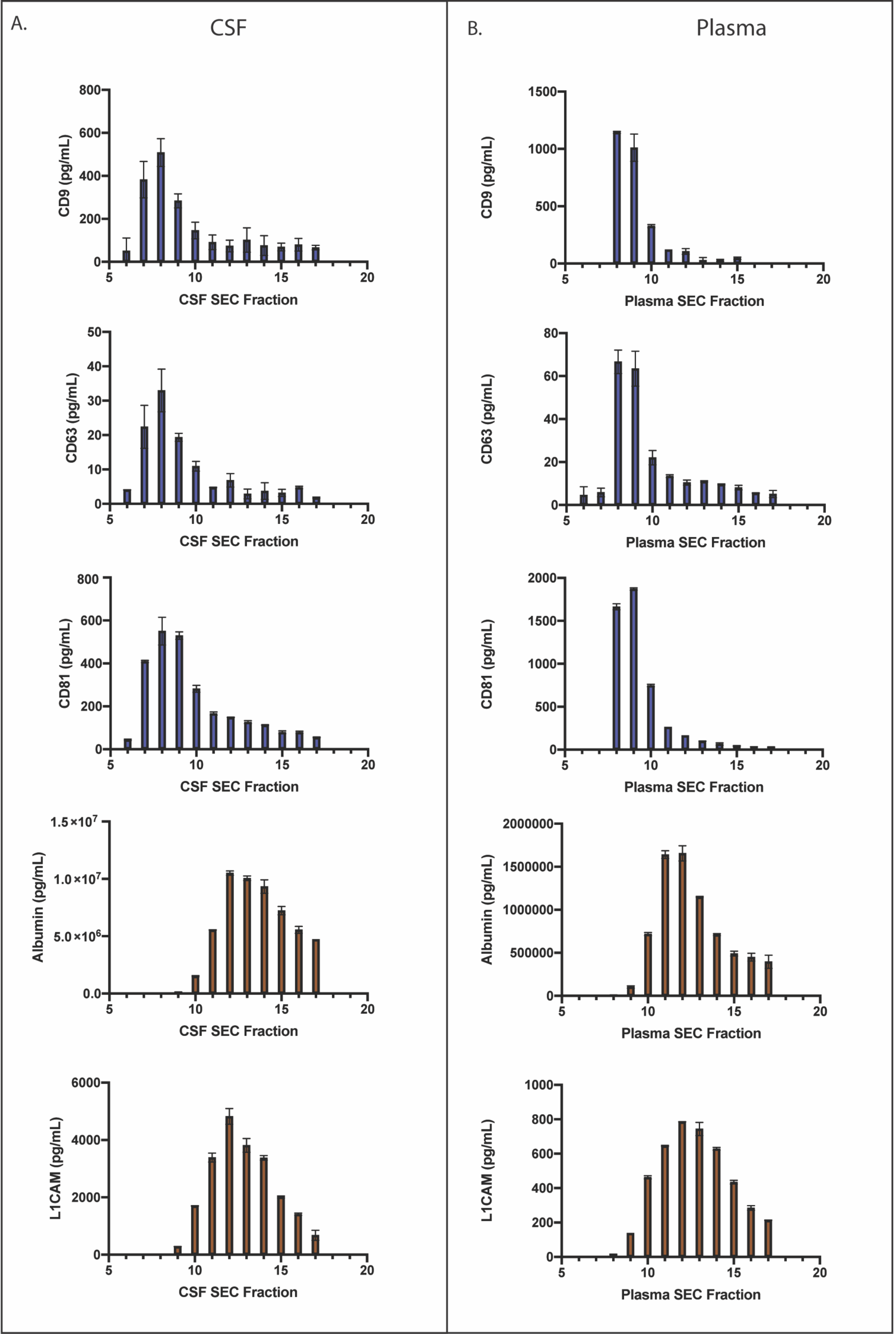
Size Exclusion Chromatography of CSF and Plasma. A) Simoa quantification of CD9, CD63, CD81, albumin and L1CAM in Sepharose 6B 10mL SEC fractions of CSF B) Simoa Quantification of CD9, CD63, CD81, albumin and L1CAM in Sepharose 6B 10mL SEC fractions of Plasma

**Figure 3:**
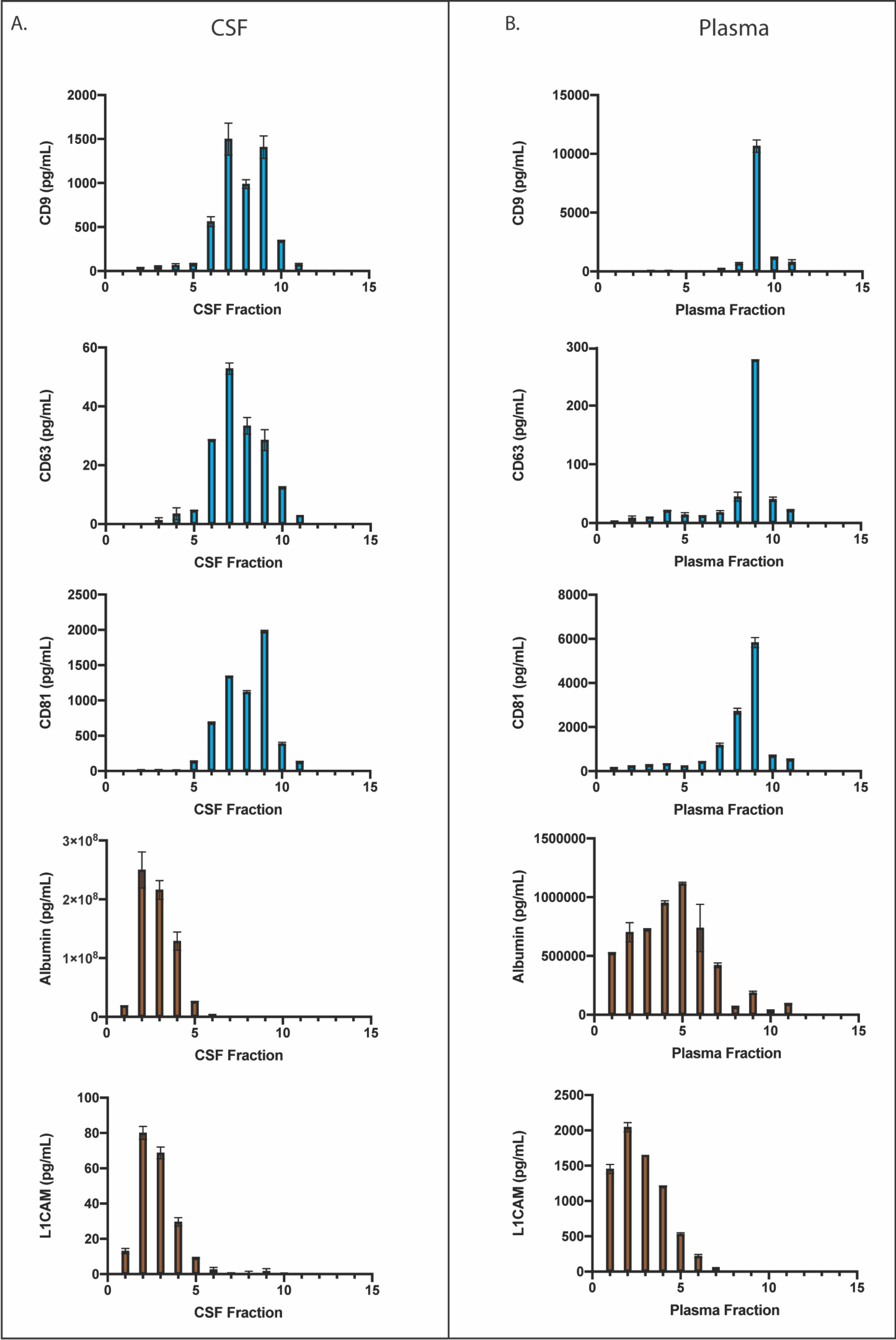
Density Gradient Centrifugation of CSF and Plasma. A) Simoa quantification of CD9, CD63, CD81, albumin and L1CAM in Density Gradient Centrifugation fractions of CSF B) Simoa Quantification of CD9, CD63, CD81, albumin and L1CAM in Density Gradient Centrifugation fractions of Plasma

### Investigation of soluble L1CAM isoforms in CSF and plasma

After observing that L1CAM appears to be in a free form and not bound to EVs in these two biofluids, we next sought to investigate the isoforms of L1CAM detected in CSF and plasma. We hypothesized two ways in which the L1CAM ectodomain detected by the Simoa assay may be present as a free protein: L1CAM present on cells may be cleaved by extracellular proteases such as ADAM10 or Plasmin [49, 50], or an isoform of L1CAM lacking a transmembrane domain may be produced by alternative splicing [48]. To investigate these possibilities, we hypothesized that proteolysis of L1CAM would produce a molecule containing the extracellular domain, but not its intracellular domain (schematically represented in Figure 4a), while alternative splicing excluding specifically the transmembrane domain would produce a molecule containing both the extracellular and intracellular domains as part of a soluble protein.

**Figure 4:**
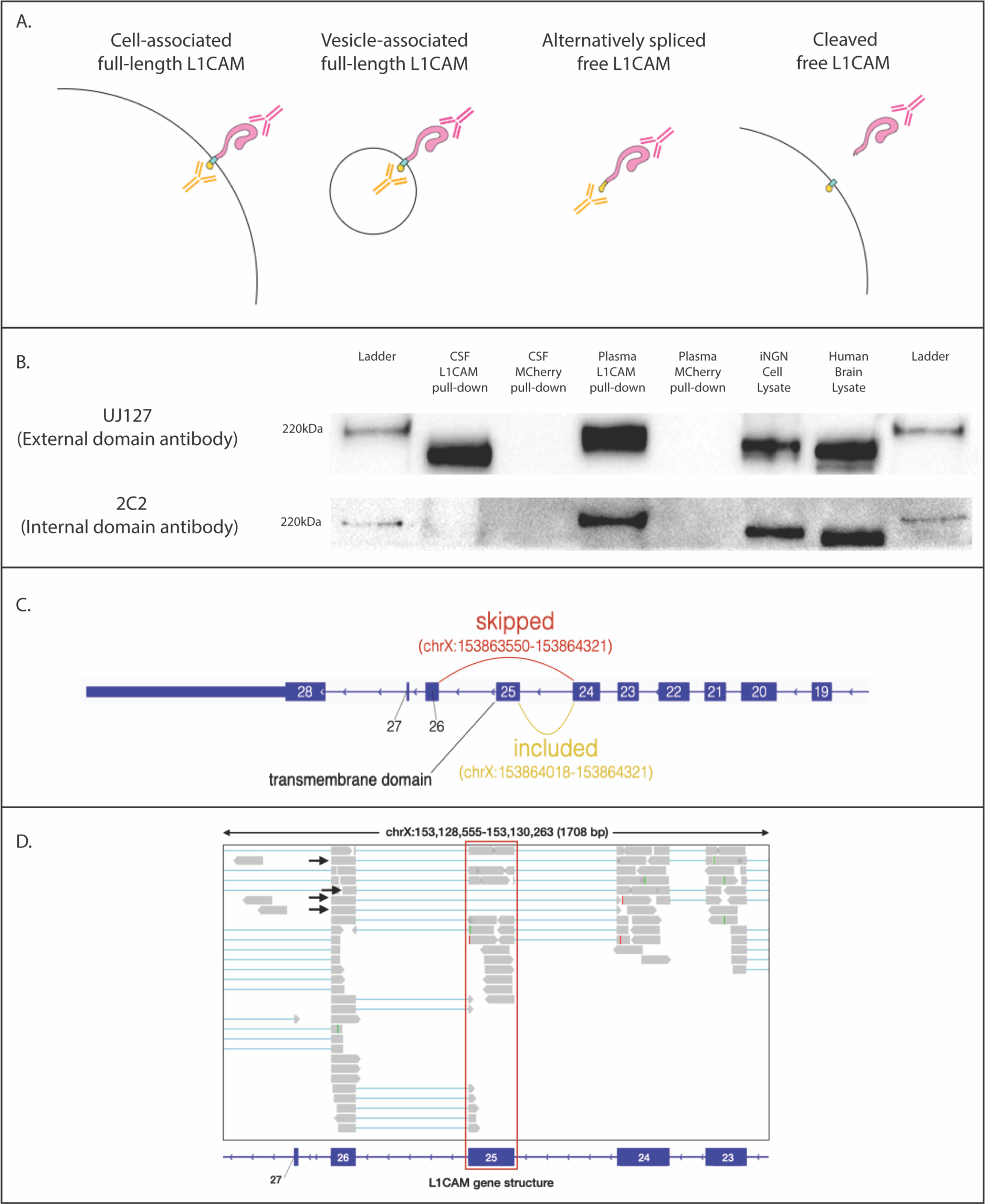
Isoforms of L1CAM in CSF and Plasma. A) Schematic representation of putative L1CAM isoforms present in the human body B) Western blotting of L1CAM immunocaptured in CSF and plasma using an antibody to the external domain of L1CAM (Clone EPR23241-224). Staining was done with one antibody to an external domain (Clone UJ127) or internal domain (Clone 2C2). C) In GTEx samples, RNA-seq supports some skipping of exon 25, as junction reads between exons 24 and 26 are detected. D) Reads from GTEx RNA-Seq data of human Tibial Artery loaded in Integrative Genome Browser (IGV) aligning to Exon 25 of L1CAM, which contain the transmembrane domain (highlighted in red). Junction reads supporting the skipping of Exon 25 are indicated with black arrows.

We immunocaptured L1CAM from both CSF and plasma and performed western blotting utilizing one antibody to the external domain and another to the internal domain. L1CAM immunocaptured from CSF produced a band at approximately 200 kDa, which blotted only with the external domain antibody, but not with the internal domain antibody. Conversely, L1CAM immunocaptured from plasma demonstrated a band at approximately 220 kDa and blotted with both the internal and external domain antibodies, (Figure 4b). Additionally, using mass spectrometry, we were able to detect a peptide matching the cytoplasmic portion of L1CAM in plasma (Supplemental Figure 3). Although we are not able to conclude whether soluble L1CAM is cleaved, alternatively spliced or both, our mass spectrometry and western blot results suggest that some proportion of soluble L1CAM in plasma is alternatively spliced to exclude the exon encoding for the transmembrane domain [48-50].

### Alternative Splicing of L1CAM Exon 25

The transmembrane domain of L1CAM is contained entirely in exon 25 (Figure 4c). Skipping of this exon leads to production of an L1CAM protein containing an ER signal sequence but no domain for insertion into the plasma membrane, one which is therefore secreted into the extracellular medium. Expressed sequence tags (ESTs) have previously been observed showing skipping of exon 25 in human endothelial cell lines [48]. We analyzed Genotype-Tissue Expression (GTEx) RNA-Seq data across diverse human organs to investigate L1CAM splicing status in other tissues. Although the junction reads supporting inclusion of exon 25 generally outnumbered those supporting skipping, we did observe reads traversing the junction from exon 24 directly to exon 26 in a number of tissues, including Tibial artery (Figure 4d). These reads in Tibial artery and other organs support the existence of an L1CAM isoform lacking its transmembrane domain and suggest that this isoform is produced at low levels across many different tissues in the human body (Supplemental Figure 4 & Supplemental Table 2).

### Antibodies to L1CAM non-specifically bind free proteins such as alpha-synuclein

Finally, we sought to understand why prior publications had found increased “cargo” proteins such as alpha-synuclein in their L1CAM EV immunocapture samples. We hypothesized that this difference may be caused by nonspecific binding of free alpha-synuclein in plasma to the L1CAM antibody used for isolation. To test this hypothesis, we followed the L1CAM immunocapture protocol of one such study utilizing the same L1CAM antibody (Clone UJ127) and nonspecific mIgG control antibodies, but we performed the isolation on recombinant alpha-synuclein instead of plasma [2]. Utilizing a Simoa assay we found that three times more recombinant alpha-synuclein binds to the UJ127 antibody than the mIgG control (Supplementary Figure 5), which matches the enrichment of alpha-synuclein found in the prior work. This result suggests that the measured changes in levels of certain proteins after EV immunocapture may be due to differences in nonspecific binding of free proteins to the capture antibodies and highlights the importance of performing rigorous controls to exclude this possibility.

## Discussion

Utilizing a variety of biochemical methods, we demonstrate that L1CAM is not associated with EVs in human CSF or plasma. Although L1CAM is canonically a surface protein in neurons, we conclude that the main forms of L1CAM present in plasma and CSF are not the transmembrane form present on EVs, but rather soluble forms, possibly generated by proteolytic cleavage or alternative splicing. We used two different fractionation techniques, SEC and DGC, to analyze the distribution of L1CAM in human plasma and CSF. By analyzing these fractions with ultrasensitive Simoa immunoassays for L1CAM along with three EV markers (CD9, CD63, CD81), and the free protein albumin, we demonstrated that L1CAM elutes in the free protein fractions and not in the EV fractions, and that this result is robustly reproducible using either fractionation technique.

Our results are consistent with a model in which the L1CAM isoform present in CSF may be produced predominantly by cleavage of L1CAM from cell surfaces, while the L1CAM isoform present in plasma arises from alternative splicing excluding exon 25 [48], but further research is needed to confirm the exact structures of L1CAM in these two sample types. Since we cannot conclusively determine the relative proportion of these isoforms, both may be present in either biofluid. Regardless of the exact isoform of L1CAM in plasma, our finding that L1CAM is not EV-associated in plasma and CSF indicates that new NDEV markers are needed. We believe that the methodology and tools we present here to investigate L1CAM can be used to validate other putative cell-type specific EV markers, not only for brain-derived EV immunocapture, but for immunocapture of EVs from any cell.

Despite this surprising result with L1CAM, we remain optimistic regarding the potential for brain derived EVs to provide a window into the brain in accessible biofluids such as plasma [53, 54]. We believe these findings simply underscore the need for carefully controlled experiments when working with these highly complex biofluids if that potential is to be realized. We are currently searching for markers other than L1CAM to isolate bona fide NDEVs from the brain using a variety of computational and experimental techniques. Using the framework presented in this paper, we plan to validate these putative markers to ensure that they are present on EVs in biofluids and are truly coming from the brain.

## Acknowledgements

The authors thank Alex Ng for help with stem cell differentiation and Jan Van Deun for help with density gradient centrifugation. The authors also thank Taplin Biological Mass Spectrometry Facility at Harvard Medical School and the Harvard Center for Mass Spectrometry for help with proteomics experiments. This work was supported by funding from the Chan Zuckerberg Initiative (CZI) Neurodegeneration Challenge Network, Good Ventures, NIH Center for Excellence in Genomic Science, Howard Hughes Medical Institute (HHMI), and the Klarman Cell Observatory (KCO).

## Disclosures

DRW is a founder and equity holder of Quanterix. AR is a SAB member of ThermoFisher Scientific, Neogene Therapeutics, Asimov and Syros Pharmaceuticals. AR is a cofounder of and equity holder in Celsius Therapeutics and an equity holder in Immunitas. From August 1, 2020, AR is an employee of Genentech. GMC is a founder, consultant, or advisory board member to companies listed here: http://arep.med.harvard.edu/gmc/tech.html

The authors have filed intellectual property on methods for EV analysis and isolation.

