## Supplementary material for "L1CAM is not Associated with Extracellular Vesicles in Human Cerebrospinal Fluid or Plasma": SI

Supplementary Information

**Supplementary Figure 1:** Linearity of dilution for Simoa assays

**Supplementary Table 1:** Spike and recovery for Simoa assays

**Supplementary Figure 2:** Comparison of ELISA and Simoa for EV quantification

**Supplementary Figure 3:** Mass spectrometry of L1CAM immunocaptured from plasma

**Supplementary Figure 4:** Analysis of reads from GTEx RNA-Seq Data indicating Exon 25 skipping in alternative splicing of L1CAM

**Supplementary Figure 5:** Affinity of L1CAM for recombinant alpha-synuclein

**Supplementary Table 2:** Analysis of reads from GTEx RNA-Seq Data supporting the existence of an L1CAM isoform lacking its transmembrane domain (table attached)

**Supplementary Figure 1:** We developed Simoa assays for the EV markers CD9, CD63, CD81 as well as for L1CAM and albumin. To ensure that our assays are accurate and precise, we performed validation experiments. Each assay demonstrated endogenous dilution linearity (parallelism) in human CSF (a) and plasma (b). Error bars represent the standard deviation from two technical replicates. Each linearity experiment was performed on the CSF and plasma from four individuals.

a.

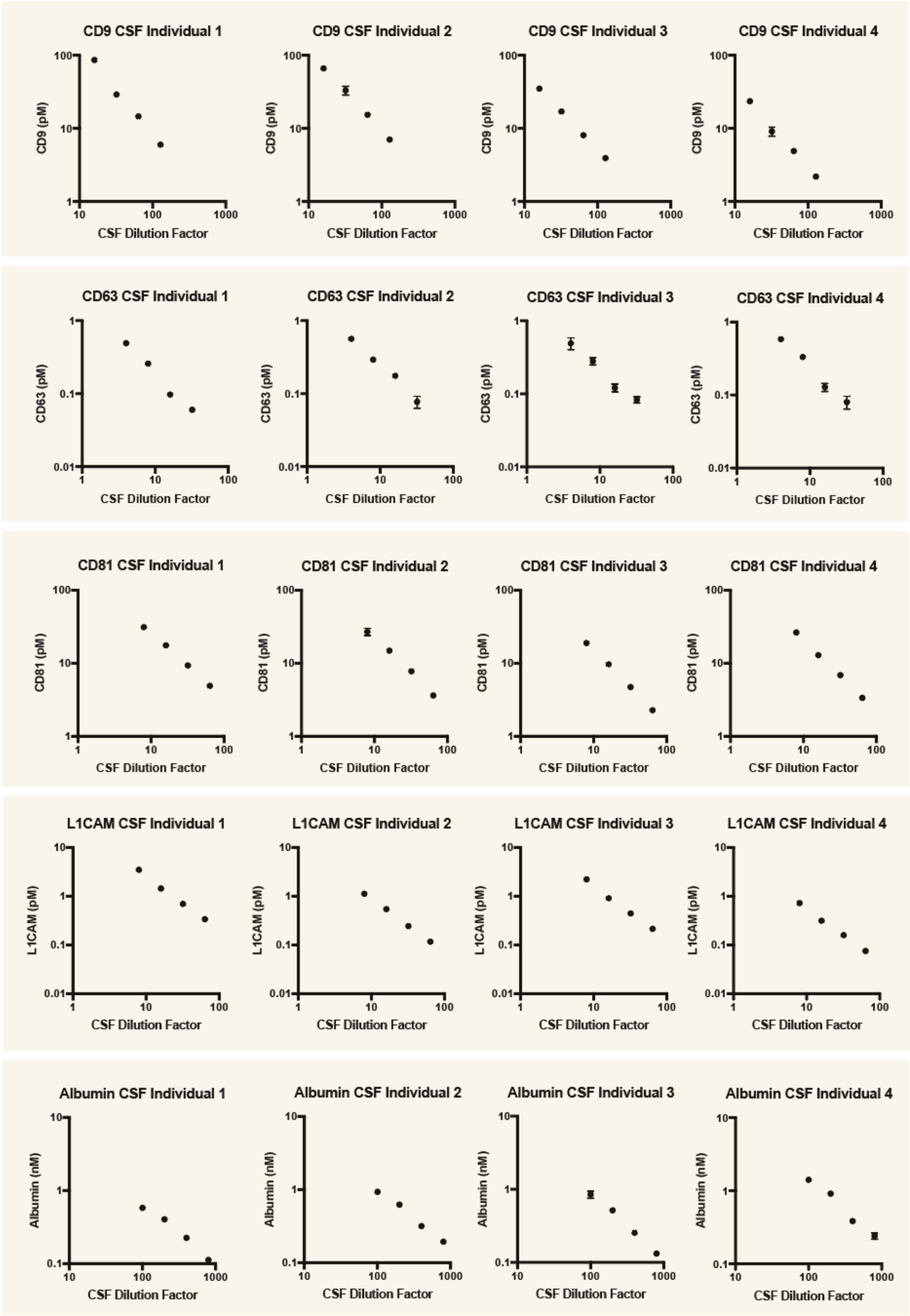

b.

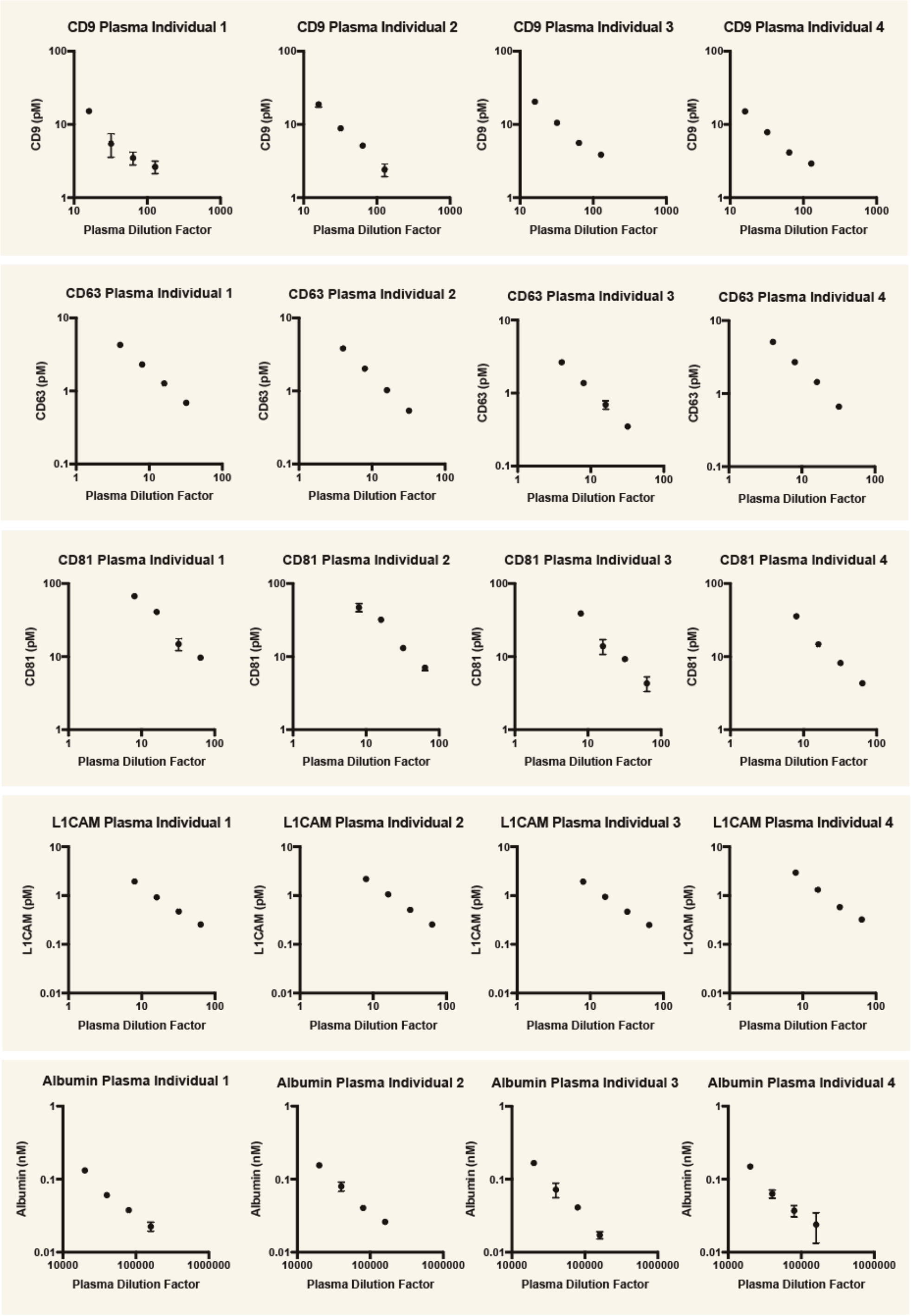
CD9 Plasma Individual 1
CD9 Plasma Individual 2
CD9 Plasma Individual 3
CD9 Plasma Individual 4
CD9 (pM)
1
10
100
1000
Plasma Dilution Factor
CD63 Plasma Individual 1
CD63 Plasma Individual 2
CD63 Plasma Individual 3
CD63 Plasma Individual 4
CD63 (pM)
0.1
Plasma Dilution Factor
CD81 Plasma Individual 1
CD81 Plasma Individual 2
CD81 Plasma Individual 3
CD81 Plasma Individual 4
CD81 (pM)
Plasma Dilution Factor
L1CAM Plasma Individual 1
L1CAM Plasma Individual 2
L1CAM Plasma Individual 3
L1CAM Plasma Individual 4
L1CAM (pM)
0.01
Plasma Dilution Factor
Albumin Plasma Individual 1
Albumin Plasma Individual 2
Albumin Plasma Individual 3
Albumin Plasma Individual 4
Albumin (nM)
10000
100000
1000000
Plasma Dilution Factor

**Supplementary Table 1:** We spiked the recombinant protein used for the calibration curve into human plasma and CSF and quantified the percent recovery. Each assay recovered between 70-130% of the spiked concentration, indicating good assay precision. Data averaged across four individuals and two technical replicates.

| Protein | CSF Dilution Factor/Spike | Average Recovery (4 Individuals) | Plasma Dilution Factor/Spike | Average Recovery (4 Individuals) |
|---|---|---|---|---|
| **CD9** | 32x+500 | 86% | 32x+500 | 79% |
| | 32x+1000 | 101% | 32x+1000 | 74% |
| **CD63** | 16x+10 | 106% | 16x+10 | 83% |
| | 16x+50 | 105% | 16x+50 | 74% |
| **CD81** | 32x+500 | 104% | 32x+500 | 106% |
| | 32x+1000 | 101% | 32x+1000 | 94% |
| **Albumin** | 400x+50000 | 90% | 80000x+50000 | 85% |
| | 400x+100000 | 85% | 80000x+100000 | 90% |
| **L1CAM** | 32x+500 | 106% | 64x+500 | 102% |
| | 32x+1000 | 107% | 64x+1000 | 98% |

**Supplementary Figure 2: Comparison of ELISA and Simoa for EV quantification**

ELISA and Simoa were used to measure tetraspanin levels. For each assay, the same sample is directly compared by ELISA and Simoa for the calibration curve using protein standard (left), EV quantification of pooled plasma SEC fractions (middle) and EV quantification of pooled CSF SEC fractions (right) (a-f). The limits of detection (LOD) of the five Simoa assays were one to two orders of magnitude more sensitive than their respective standard ELISAs using the same set of antibodies for each target. Error bars represent the standard deviation from two technical replicates.

To ascertain whether this improved sensitivity was necessary for quantifying EVs in biological samples, human plasma and CSF samples were fractionated using a commercially available SEC column (Izon qEVoriginal 35nm), and fractions were evenly split for downstream analysis using Simoa, ELISA, and western blotting. Simoa quantified tetraspanins in EV fractions 7-10 (early fractions where EVs are expected) for all markers in both plasma and CSF (b,d,f). ELISA was able to quantify CD9 in expected EV fractions (7-10) for plasma, while only a subset of the expected EV fractions were quantifiable in plasma for CD63 and CD81. In contrast, none of the CSF fractions 7-10 were quantifiable by ELISA for any of the tetraspanins (a,c,e). When the SEC fractions were analyzed by western blot using the same volume of each fraction, we observed very high background in the plasma fractions (due to the high abundance of free proteins), but the tetraspanins were detectable and matched the general pattern of the Simoa results. We did not detect any tetraspanins in the CSF fractions (g). Simoa was the only assay method that allowed quantification of EVs in SEC fractions for both plasma and CSF.

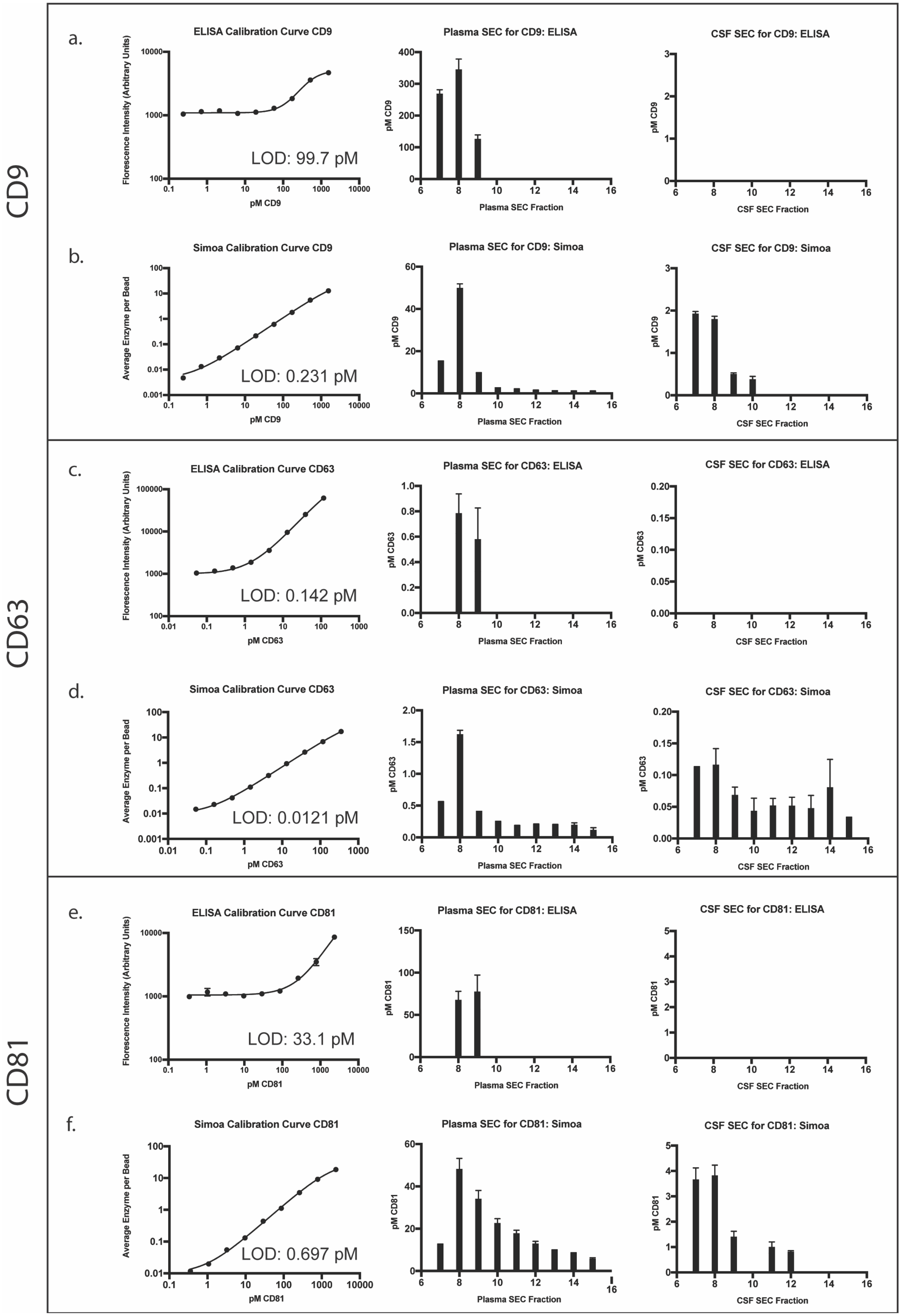
CD9
a.
ELISA Calibration Curve CD9
Florescence Intensity (Arbitrary Units)
pM CD9
LOD: 99.7 pM
Plasma SEC for CD9: ELISA
pM CD9
Plasma SEC Fraction
CSF SEC for CD9: ELISA
pM CD9
CSF SEC Fraction
b.
Simoa Calibration Curve CD9
Average Enzyme per Bead
pM CD9
LOD: 0.231 pM
Plasma SEC for CD9: Simoa
pM CD9
Plasma SEC Fraction
CSF SEC for CD9: Simoa
pM CD9
CSF SEC Fraction
CD63
c.
ELISA Calibration Curve CD63
Florescence Intensity (Arbitrary Units)
pM CD63
LOD: 0.142 pM
Plasma SEC for CD63: ELISA
pM CD63
Plasma SEC Fraction
CSF SEC for CD63: ELISA
pM CD63
CSF SEC Fraction
d.
Simoa Calibration Curve CD63
Average Enzyme per Bead
pM CD63
LOD: 0.0121 pM
Plasma SEC for CD63: Simoa
pM CD63
Plasma SEC Fraction
CSF SEC for CD63: Simoa
pM CD63
CSF SEC Fraction
CD81
e.
ELISA Calibration Curve CD81
Florescence Intensity (Arbitrary Units)
pM CD81
LOD: 33.1 pM
Plasma SEC for CD81: ELISA
pM CD81
Plasma SEC Fraction
CSF SEC for CD81: ELISA
pM CD81
CSF SEC Fraction
f.
Simoa Calibration Curve CD81
Average Enzyme per Bead
pM CD81
LOD: 0.697 pM
Plasma SEC for CD81: Simoa
pM CD81
Plasma SEC Fraction
CSF SEC for CD81: Simoa
pM CD81
CSF SEC Fraction

g.

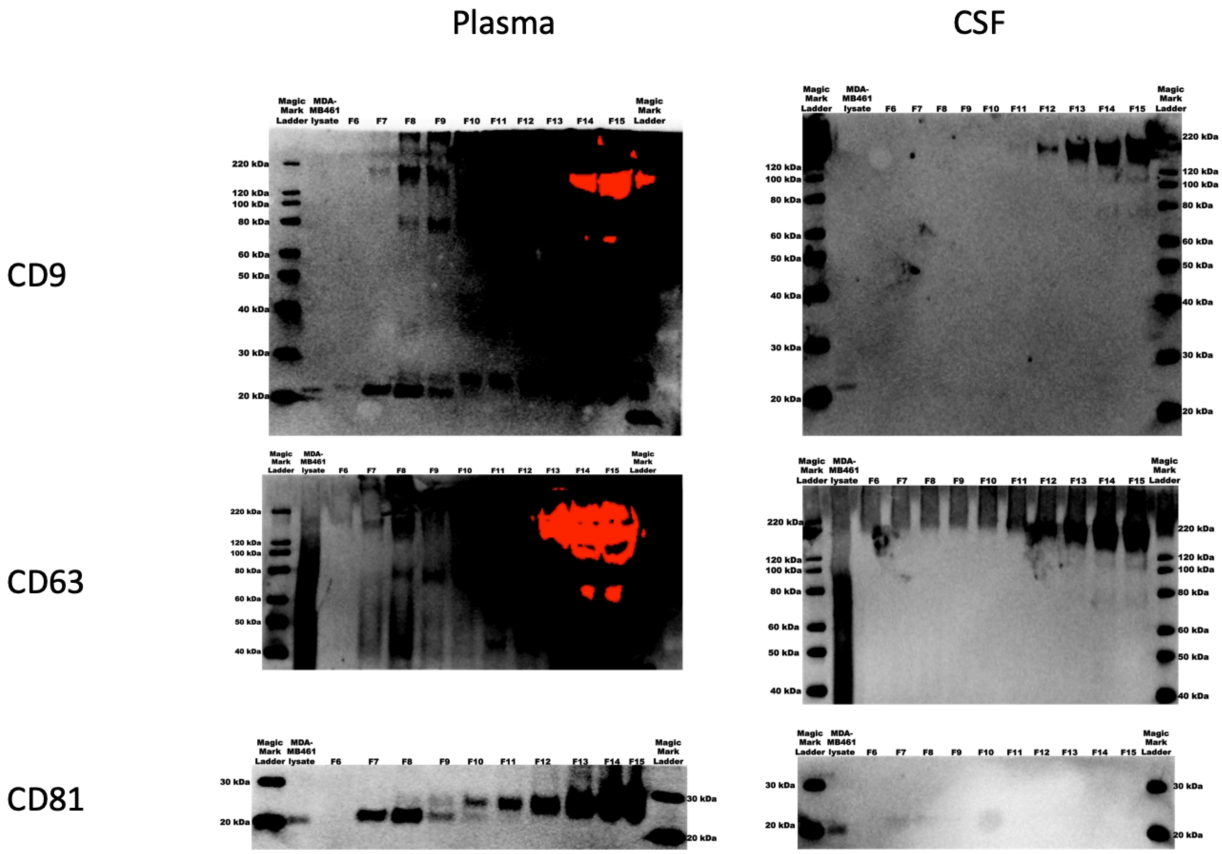

Plasma
CSF
CD9
CD63
CD81
Magic Mark Ladder
MDA-MB461 lysate
F6
F7
F8
F9
F10
F11
F12
F13
F14
F15
Magic Mark Ladder
220 kDa
120 kDa
100 kDa
80 kDa
60 kDa
50 kDa
40 kDa
30 kDa
20 kDa

**Supplementary Figure 3: Mass spectrometry of L1CAM immunocaptured from plasma**
Mass spectrometry of full length recombinant L1CAM protein standard shows, as expected, peptides matching an isoform which includes Exon 25 (a). Mass Spectrometry of L1CAM immunocaptured from human plasma shows peptides matching the cytosolic domain at the C terminus (emphasized with black arrow) (b). Full length sequence of L1CAM displayed with peptides from mass spectrometry shown in green. Blue box indicates L1CAM transmembrane domain and red box indicates amino acid sequence encoded by Exon 25

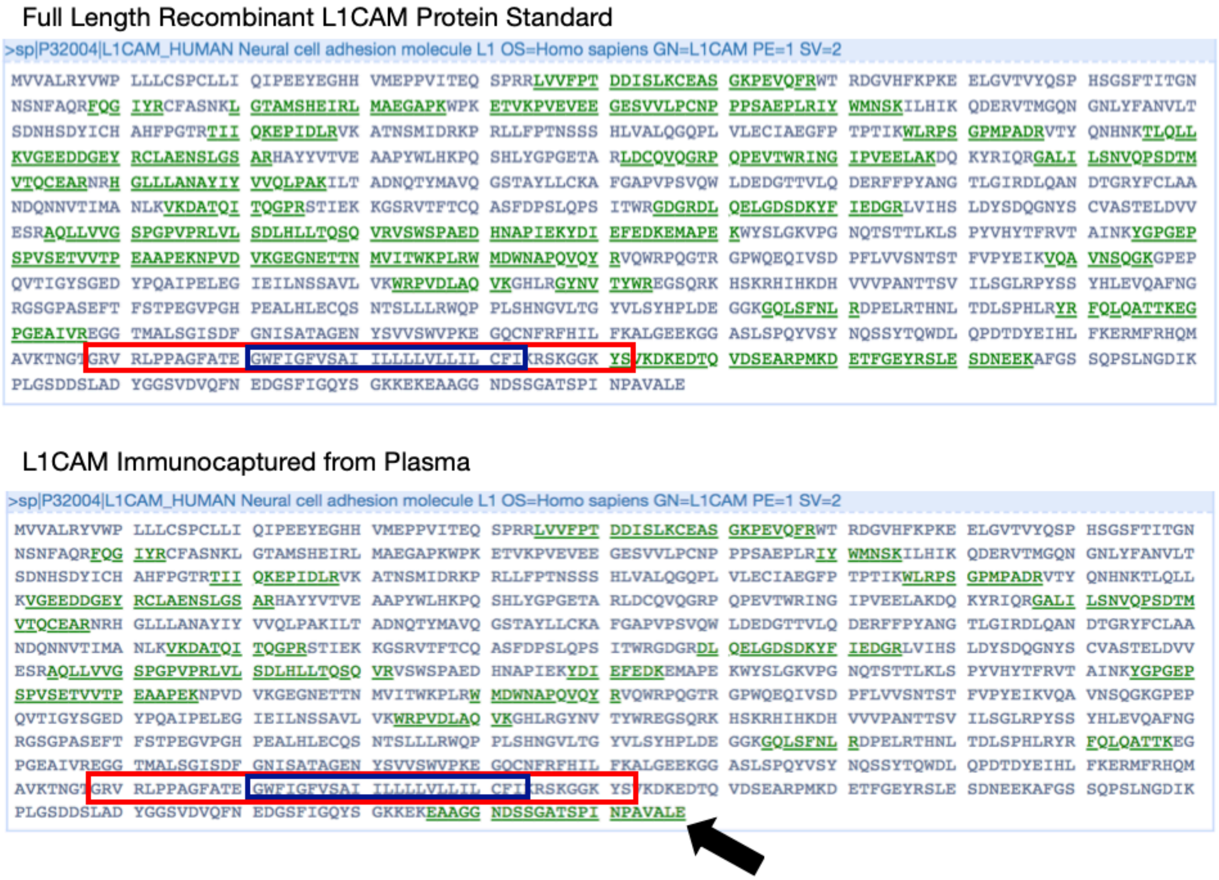

**Supplementary Figure 4: Analysis of reads from GTEx RNA-Seq Data indicating Exon 25 skipping in alternative splicing of L1CAM**

Fraction of reads mapping to L1CAM isoform supporting skipping of L1CAM Exon 25 (junction reads spanning Exon 24 and Exon 26) vs. inclusion of Exon 25 from RNA-Seq GTEx data of various human organs.

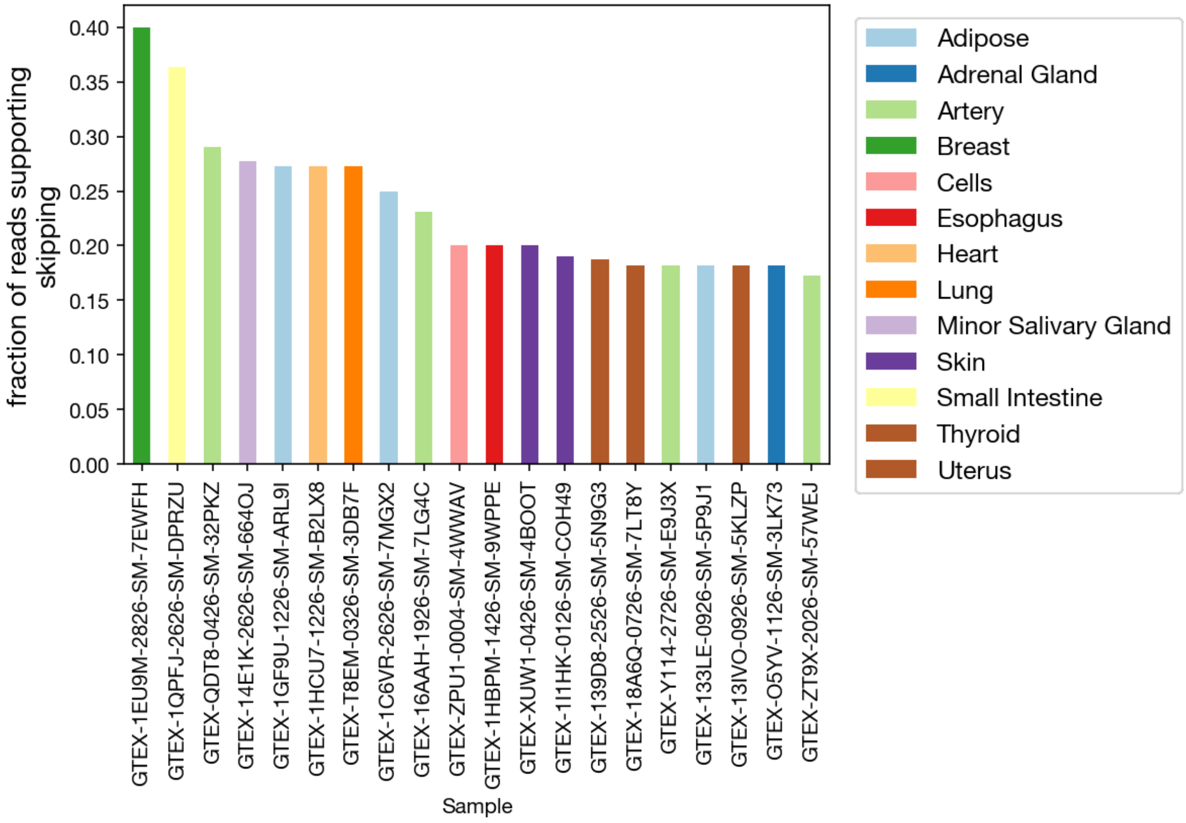

**Supplementary Figure 5: Affinity of L1CAM for recombinant alpha-synuclein**

In order to assess nonspecific binding of UJ127 to alpha-synuclein, we followed the protocol from Min Shi et al. 2014. We utilized the same L1CAM antibody (Clone UJ127, Abcam) and nonspecific mIgG control antibodies (Santa Cruz Biotechnology) but used recombinant alpha-synuclein instead of plasma [1]. We found that that three times more recombinant alpha-synuclein binds to the UJ127 antibody than the mIgG control, exactly mirroring the effect in the paper.

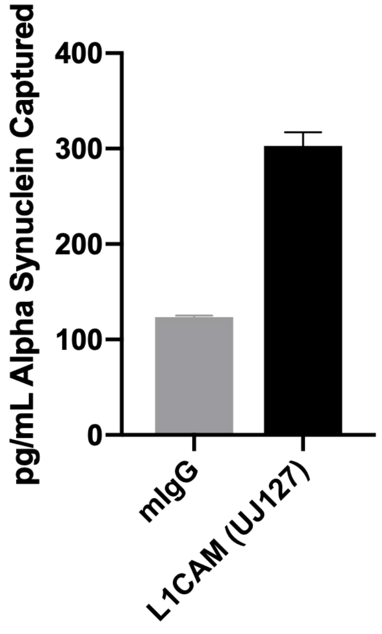



1. Shi, M., et al., *Plasma exosomal alpha-synuclein is likely CNS-derived and increased in Parkinson's disease.* Acta Neuropathol, 2014. **128**(5): p. 639-650.
