## Supplementary material for "L1CAM is not Associated with Extracellular Vesicles in Human Cerebrospinal Fluid or Plasma": Methods

Materials & Methods

**Human Sample Handling**

Pre-aliquoted human plasma and CSF samples were ordered from BioIVT. For spike and recovery and dilution linearity experiments, individual CSF and plasma from BioIVT were utilized. For all other experiments, pooled CSF and plasma were used. Plasma or CSF was thawed at room temperature. Immediately after the sample had thawed, 100X Protease/Phosphatase Inhibitor Cocktail (Cell Signaling Technology) was added at 1X final concentration to the sample. The sample was then centrifuged at 2000 x *g* for 10 minutes. Next, the supernatant was centrifuged through a 0.45μm Corning Costar SPIN-X centrifuge tube filter (Sigma-Aldrich) at 2000 x *g* for 10 minutes to get rid of any remaining cells or cell debris.

**Simoa Assays**

Candidate capture antibodies were coupled to carboxylated paramagnetic beads from the Simoa Homebrew Assay Development Kit (Quanterix) using EDC chemistry (Thermo Fisher Scientific). Candidate detection antibodies were conjugated to biotin using EZ-Link NHS-PEG4 Biotin (Thermo Fisher Scientific). Reagents were cross-tested for signal against the following recombinant proteins for CD9, CD63, CD81, L1CAM and albumin: ab152262 (Abcam), TP301733 (Origene), CD81 Origene TP317508 (Origene), TP311601 (Origene), ab201876 (Abcam) on a Simoa HD-X Analyzer (Quanterix). The antibody pairs that gave the highest signal to background ratio were further validated in two ways: First, plasma and CSF were serially diluted in sample buffer in order to demonstrate endogenous dilution linearity (Supplementary Information Figure 1). Next, the recombinant protein used in the calibration curve was added to plasma and CSF at two different concentrations in order to determine that spike and recovery was within 70-130% (Supplementary Information Table 1).

For CD9, CD63, CD81 and L1CAM assays, samples were incubated with immunocapture beads (25 μL) and biotinylated detection antibody (20 μL) for 35 minutes. Next, six washes were performed, and the beads were resuspended in 100 μL of Streptavidin labeled β-Galactosidase (Quanterix) and incubated for 5 minutes. An additional six washes were performed, and the beads were resuspended in 25 μL Resorufin β-D-Galactopyranoside (Quanterix) before being loaded into the microwell array.

For the albumin assay, samples were incubated first with immunocapture beads (25 μL) for 15 minutes and then washed six times. Subsequently, 100 μL detection antibody was incubated with the beads for 5 minutes. Next, six washes were performed, and the beads were resuspended in 100 μL of Streptavidin labeled β-Galactosidase (Quanterix) for a final 5-minute incubation. An additional six washes were performed, and the beads were resuspended in 25 μL Resorufin β-D-Galactopyranoside (Quanterix) before being loaded into the microwell array.

**Plate-Based ELISA**

Simoa assays were transferred to a plate-based ELISA for comparison. Capture antibody was diluted in ELISA Coating Buffer (BioLegend) at a concentration of 4 μg/mL and 100 μL was coated per well on a Nunc MaxiSorp ELISA plate (BioLegend). Plates were incubated with capture antibody overnight at 4 °C. Subsequently, the plate was washed 3 times with 200 μL PBST. Sample was added to each well and incubated at room temperature for 3 hours. The plate was again washed 3 times with 200 μL PBST. 100 μL of corresponding detection antibody was added to the plate and left to incubate for 1 hour. Detection antibody was then removed, and the plate was washed 3 times with 200 μL PBST. 100 μL Streptavidin labeled β-Galactosidase from the Simoa Homebrew Assay Development Kit (Quanterix) was then added and incubated for 30 minutes. The plate was then washed 5 times with 200 μL PBST and incubated with 100 μL of Resorufin β-D-Galactopyranoside, also from the Simoa Homebrew Assay Development Kit (Quanterix), for 20 minutes in the dark. Plates were then imaged with a Tecan Plate Reader using Magellan v 7.2 software at 555 nm excitation and 605 nm emission.

**Western Blotting**

Western blotting was performed as previously described in detail [1].

*Western blot for Supplementary Figure 2:*

Equal volumes of each SEC fraction were loaded on a Bolt 4-12% Bis-Tris Plus gel (Thermo Fisher Scientific) after addition of Bolt LDS sample buffer (Thermo Fisher Scientific) and denaturation at 70°C for 10 minutes. The gel was run at 150 V for 60 minutes and then transferred onto a nitrocellulose membrane using the iBlot2 Dry Blotting System (Thermo Fisher Scientific). The following primary antibodies were used for western blot at the corresponding dilutions in milk overnight at 4°C: MM2/57 for CD9 (Millipore Sigma) at 1:1000, h5c6 for CD63 (BD Biosciences) at 1:1000, M38 for CD81 (Thermo Fisher Scientific) at 1:666. After three washes with PBST, anti-mouse IgG Secondary antibody (Bethyl Laboratories) was added for 2 hours in milk buffer at 1:2000. After three more washes in PBST, SpectraQuant HRP-CL Spray Chemiluminescent Detection Reagent (BridgePath Scientific) was used to develop the western blots. Imaging was performed on the Sapphire Biomolecular Imager (Azure Biosystems).

*Immunocapture & Western blot for Figure 4b:*

Immunocapture of L1CAM from plasma and CSF was performed using the Dynabeads antibody coupling kit (Invitrogen 14311D). Following the manufacturer's instructions either anti-L1CAM antibody (Abcam ab272321) or anti-MCherry control antibody (Abcam ab232341). Four aliquots of 1mL of CSF or .375mL of plasma were incubated at 4°C overnight with gentle rotation. Plasma or CSF were washed 4 times with PBS and resuspended in LDS with reducing buffer. Immunocaptured plasma and CSF L1CAM as well as human brain lysate and iNGN cell lysate were loaded on a Bolt 4-12% Bis-Tris Plus gel (Thermo Fisher Scientific) after addition of Bolt LDS sample buffer with reducing buffer (Thermo Fisher Scientific) and denaturation at 70°C for 10 minutes. The gel was run at 150V for 120 minutes and then transferred onto a nitrocellulose membrane using the iBlot2 Dry Blotting System (Thermo Fisher Scientific). The following primary antibodies were used for western blot at the corresponding dilutions in milk overnight at 4°C: UJ127 (Abcam) at 1:2000 and 2c2 (Abcam) at 1:500. After three washes with PBST, anti-mouse cross IgG secondary antibody (Bethyl Laboratories) was added for 2 hours in milk buffer at 1:2000. After three more washes in PBST, SpectraQuant HRP-CL Spray Chemiluminescent Detection Reagent (BridgePath Scientific) was used to develop the western blots. Imaging was performed on the Sapphire Biomolecular Imager (Azure Biosystems).

**Preparation of Custom SEC Columns**

Sepharose CL-6B (GE Healthcare) was washed in PBS. Briefly, the volume of resin was washed with an equal volume of PBS in a glass jar and then placed at 4 °C in order to let the resin settle completely (several hours or overnight). The PBS was then poured off, and new PBS was again added for a total of 3 washes. Columns were prepared fresh on the day of use. Washed resin was poured into an Econo-Pac Chromatography column (Bio-Rad) for a 10 mL bed volume. The column was allowed to drip out until the column was solid at which time the top frit was placed securely at the top of the resin but without compression. PBS was then added at 1 mL above the frit until sample was ready to be added.

**Collection of Size Exclusion Chromatography Fractions**

Once prepared, all columns were washed with at least 20 mL of PBS in the column. When sample was ready to be loaded, the column was allowed to fully drip out and, after last drop, plasma or CSF was added to the column. Immediately thereafter, 0.5 mL fractions were collected. As soon as the plasma or CSF completely went through the frit, PBS was added to top of column 1 mL at a time. For plasma and CSF, fractions 6-19 were collected while for iNGN fractions 6-15 were collected. For the comparison of Simoa, ELISA, and western blot, one 0.5 mL sample was fractionated by SEC using an Izon qEVoriginal 35nm column into 0.5 mL fractions (fractions 6-15). Each of the fractions was then divided evenly for use in the three techniques.

**Density Gradient**

For the density gradient centrifugation, an Optiprep (Iodixanol) gradient was prepared using the following layers (from top to bottom): 2 mL 5%, 3mL 10%, 3mL 20%, 3 mL 40% layers. Each layer of Optiprep (Millipore Sigma) was diluted in solution of 0.25M Sucrose (Millipore Sigma) and Tris-EDTA pH 7.4 (Millipore Sigma) . Sample (1mL) was loaded on top of the 5% fraction. Samples were centrifuged at 100,000 RCF for 18 hours at 4C in 13.2 Polypropylene tubes (Beckman Coulter). Fractions were removed from the top in 1 mL increments.

**Mass Spectrometry:**

Excised gel bands around the size of 180-240kDa were cut from a polyacrylamide gel and submitted for mass spectrometry analysis. Gel pieces were washed and dehydrated with acetonitrile, and then acetonitrile was removed. Gel pieces were dried in Speed-Vac and a modified in-gel Trypsin procedure was performed [2]. Samples were then reconstituted in 5 - 10 µl of HPLC solvent A (2.5% acetonitrile, 0.1% formic acid). A nano-scale reverse-phase HPLC capillary column was created by packing 2.6 µm C18 spherical silica beads into a fused silica capillary with a flame-drawn tip [3]. Each sample was loaded via a Famos auto sampler (LC Packings) onto the column. Peptides were eluted with increasing concentrations of solvent B

(97.5% acetonitrile, 0.1% formic acid), subjected to electrospray ionization, and entered into an LTQ Orbitrap Velos Pro ion-trap mass spectrometer (Thermo Fisher Scientific). Peptide sequences were determined by matching protein databases with the acquired fragmentation pattern by the software program, Sequest (Thermo Fisher Scientific) [4].

**iNGN EV isolation:**

Previously described iNGN cells were grown in mTeSR1 media (STEMCELL Technologies) on Matrigel (Corning) coated plates. Doxycycline (Sigma Aldrich) was diluted in PBS and added to MTeSR1 at a final concentration of 0.5 µg/mL to initiate differentiation. On Day 4 after Dox addition, media was switched to Gibco DMEM with Glutamax (Thermo Fisher Scientific) supplemented with B27 Serum-Free Supplement (Thermo Fisher Scientific) and Gibco Penicillin Streptomycin (Thermo Fisher Scientific). On Day 6, EVs were isolated by differential ultracentrifugation, as previously described [5], and resuspended in PBS.

**Splicing RNA-Seq Analysis**

For analysis of Genotype-Tissue Expression database (GTEx) data, pre-processed exon-exon junction read- counts were obtained from the publicly available version 8 database (https://GTExportal.org). The read table was filtered to include only junctions falling within the body of L1CAM and these counts were analyzed using custom python scripts available upon request. Reads aligning to L1CAM (chrX:153,859,517-153,888,173) were visually examined using the Integrative Genomics Viewer (IGV) [6] to assess evidence of inclusion or exclusion of exon 25.
